# Loss of SEZ6L2 disrupts synaptic architecture and drives progressive behavioral deficits

**DOI:** 10.64898/2025.12.31.697214

**Authors:** Julia M. Granato, Nina Silver, Alison Hobbins, John Randolph, Angela Stout, Harris A. Gelbard, Jenny M. Gunnersen, Samuel J. Mackenzie, Patricia M. White, Jennetta W. Hammond

**Affiliations:** University of Rochester Medical Center, Rochester, NY; University of Melbourne, Melbourne, VIC 3010, Australia

**Author notes:** **Correspondence to:** Jennetta W. Hammond, 601 Elmwood Ave, Box 645, Rochester, NY 14526.

**Keywords:** SEZ6, SEZ6L, SEZ6L2, behavior, cortex, neuron, motor coordination, 16p11

## Abstract

The SEZ6 family, composed of SEZ6, SEZ6L, and SEZ6L2, plays essential roles in neurodevelopment, synaptic maintenance, and complement regulation. However, the unique contribution of SEZ6L2 to brain development and function remains largely unexplored. In this study, we provide the first comprehensive behavioral and neurobiological characterization of *Sez6l2* knockout (KO) mice and directly compare their phenotype with *Sez6* triple knockout (TKO) mice, which lack all three *Sez6* family genes. *Sez6l2* KO mice exhibit impairments across multiple behavioral domains, including motor coordination, gait, sociability, sensory processing, and goal-directed repetitive behaviors. Several phenotypes, particularly motor deficits, worsen with age. Male *Sez6l2* KO mice also demonstrate enhanced fear learning and increased prepulse inhibition, revealing sex-specific alterations in sensorimotor gating. At the synaptic level, *Sez6l2* KO mice show reduced dendritic spine length and decreased expression of key postsynaptic proteins consistent with impaired excitatory synaptic connectivity. In comparison, *Sez6* TKO mice display more severe impairments across most measures. To examine the contribution of complement signaling, complement component C3 was deleted in *Sez6* TKO mice. This manipulation rescued dendritic spine deficits and partially improved select gait parameters but did not restore broader behavioral outcomes. These findings indicate that complement dependent mechanisms contribute to specific structural and motor phenotypes but are not sufficient to explain the full behavioral deficits. SEZ6L2 therefore emerges as a critical regulator of synaptic integrity and behavior, providing a framework for understanding how SEZ6 family dysfunction may contribute to neurodevelopmental and neurodegenerative disorders.

## INTRODUCTION

The SEZ6 family, comprising SEZ6, SEZ6L, and SEZ6L2, is associated with several neurological disorders, including autism spectrum disorder (ASD) (Kumar et al. 2009; Konyukh et al. 2011; Chapman et al. 2015; Lim et al. 2017; Zhou et al. 2022; Kim et al. 2024), schizophrenia (Ambalavanan et al. 2016), bipolar disorder (Xu et al. 2013), intellectual disability (Gilissen et al. 2014), autoimmune cerebellar ataxia (Yaguchi et al. 2014; Landa et al. 2021; Abe et al. 2023), and Alzheimer’s disease (AD) (Paracchini et al. 2018; Nguyen et al. 2024; Cheng et al. 2025). Notably, *Sez6l2* is part of the 16p11.2 CNV region, which accounts for ∼1% of ASD diagnoses and is also strongly linked to schizophrenia (Weiss et al. 2008; Kumar et al. 2009; McCarthy et al. 2009).

SEZ6 family proteins are primarily expressed by neurons and neuroendocrine cells, with all neuronal subtypes expressing at least one member. During early development, SEZ6 proteins are widely expressed. In adults, SEZ6 is restricted to specific brain regions (e.g., deep cortical layers, striatum, hippocampal CA1), while SEZ6L and SEZ6L2 maintain broader expression throughout the brain (Kim et al. 2002; Miyazaki et al. 2006; Gunnersen et al. 2009; Osaki et al. 2011; Pigoni et al. 2016; Nash et al. 2020).

The SEZ6 family are transmembrane proteins containing three CUB and five CCP domains (Miyazaki et al. 2006). These domains are commonly found in proteins regulating the complement pathway. In the brain, this pathway drives synaptic pruning during neurodevelopment and neurodegenerative disease (Stevens et al. 2007; Kolev et al. 2009; Gonzalez-Calvo et al. 2022). Interestingly, SEZ6L2 has been shown to inhibit the complement pathway by facilitating the cleavage and decay of C3 convertases (Qiu et al. 2021). This implicates SEZ6L2 as a neuronal protection factor against excessive complement activity. Proteins that contain CCP or CUB domains also contribute to processes such as neuronal migration, dendrite outgrowth, protein trafficking, and regulation of glutamatergic receptors (Tiao et al. 2008; Ng et al. 2009; Tang et al. 2011; Salmi et al. 2013; Gonzalez-Calvo et al. 2021). Consistent with these roles, SEZ6 and SEZ6L2 interact with kainate and AMPA receptors, respectively. These interactions regulate receptor post-translational modification and surface expression (Yaguchi et al. 2017; Pigoni et al. 2020). SEZ6 proteins also aid in trafficking neuronal proteins like cathepsin D and motopsin (Mitsui et al. 2013; Boonen et al. 2016).

Knockout (KO) studies link SEZ6 proteins to synapse development and function. *Sez6* KO mice show increased dendritic branching, reduced spine density, and altered excitatory post-synaptic potentials (EPSPs) (Gunnersen et al. 2007; Zhu et al. 2018). *Sez6* TKO mice exhibit lower spine density, abnormal Purkinje cell innervation, and impaired neuronal communication (Miyazaki et al. 2006; Nash et al. 2020). Furthermore, RNAi knockdown of SEZ6 family members in neuronal cultures reduces the frequency and amplitude of Ca^2+^ transients (Anderson et al. 2012).

Synaptic alterations may contribute to the mild behavioral abnormalities reported in individual knockout models of *Sez6* and *Sez6l* (Miyazaki et al. 2006; Gunnersen et al. 2007; Ong-Palsson et al. 2022). In contrast, triple knockout of all three SEZ6 proteins results in significant motor deficits, impaired learning and memory, and increased anxiety-like behavior (Miyazaki et al. 2006; Nash et al. 2020). This raises the possibility of functional redundancy in SEZ6 family proteins and underscores the need to study both the individual and combined roles of SEZ6 proteins.

The behavioral and neurological phenotype of *Sez6l2* KO mice remains uncharacterized, limiting insight into *Sez6l2* function and its contribution to the *Sez6* TKO phenotype. Here, we assess behavioral, synaptic, and molecular consequences of *Sez6l2* loss and directly compare *Sez6l2* KO, *Sez6* TKO, and wild-type (WT) mice to define shared and nonredundant roles within the SEZ6 family. We also examine the contribution of complement signaling by testing whether genetic deletion of complement component C3 modifies synaptic and behavioral phenotypes in *Sez6* TKO mice.

## RESULTS

### *Sez6l2* KO and *Sez6* TKO mice show impaired motor functions that worsen with age

Mobility was evaluated in WT (C57BL/6), *Sez6l2* KO, and *Sez6* TKO mice using open field and gait analysis assays at 2.5 or 4 months of age, respectively. In the open field assay, female *Sez6l2* KO mice exhibited an intermediate phenotype between WT and *Sez6* TKO mice, with shorter travel distances, increased immobility, and slower maximum speeds (Fig. 1A-D). Male *Sez6l2* KO mice displayed similar impairments, while male *Sez6* TKO mice traveled distances comparable to WT mice, but still had reduced maximum speeds. Interestingly, both male and female *Sez6l2* KO mice spent more time in the center zone, suggesting decreased anxiety, while female *Sez6* TKO mice spent less time in the center, suggesting heightened anxiety in addition to their motor deficits.

**Figure 1:**
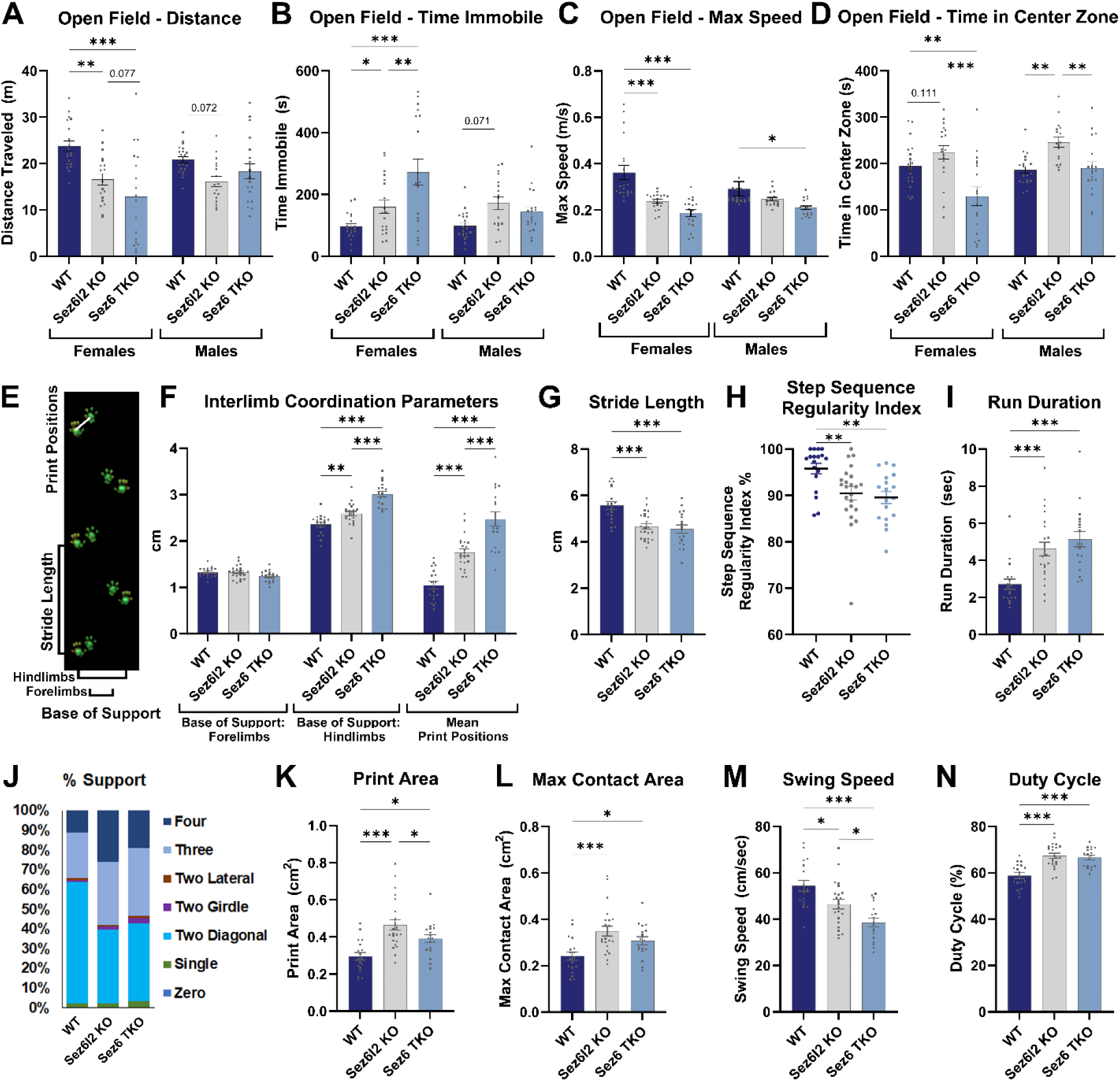
*Sez6l2* KO and *Sez6* TKO mice display impaired mobility and an altered gait at 2-4 months. (**A-D)** WT, *Sez6l2* KO, and *Sez6* TKO mice were assessed in an open field test for **A)** distance traveled, **B)** time immobile, **C)** maximum speed, and **D)** time spent in the center zone. Statistics for A-D: 2-way ANOVAs with HS MCTs; n=18-20F, 17-20M. **E)** Representative paw print image. (**F-N)** Gait analysis metrics: **F)** interlimb coordination parameters, **G)** stride length, **H)** step sequence regularity index, **I)** run duration, **J)** percent support, **K)** paw print area, **L)** paw max contact area, **M)** limb swing speed, and **N)** limb duty cycle. Statistics for E, G, J-M: 1-way ANOVAs with HS MCTs; for F: 2-way ANOVA with HS MCTs; for H: Welch ANOVA with D.T3 MCTs. For all gait metrics. n=18-23.

Gait analysis showed *Sez6l2* KO mice moved more slowly than WT and had a wider hind limb stance and increased print position distances, though less pronounced than in *Sez6* TKO mice (Fig. 1E,F). Both *Sez6l2* KO and *Sez6* TKO mice exhibited shorter stride lengths, fewer regular step sequences, shorter run durations, and altered limb support distributions (Fig. 1G-J). Across all limbs, both KO strains had larger paw prints, reduced swing speeds, and increased duty cycles (percent of step cycle with paw contact) (Fig. 1I-N). Overall, *Sez6l2* KO mice displayed impaired gait with metrics either comparable to *Sez6* TKO mice or intermediate between WT and *Sez6* TKO phenotypes.

Given the age-related motor-decline reported in *Sez6* TKO mice (Nash et al. 2020), we investigated whether *Sez6l2* KO mice exhibit similar patterns of age-dependent deterioration. At 3 months, *Sez6l2* KO performed comparably to WT mice, on the fixed-speed rotarod while *Sez6* TKO fell off significantly earlier (Fig. 2A). By 12 months, *Sez6l2* KO performance declined to an intermediate level between WT and *Sez6* TKO mice. On the accelerating rotarod, *Sez6l2* KO mice showed poorer performance than WT starting at 6 months and failed to demonstrate significant learning across trials, unlike WT mice, which showed improvement at 6 and 8 months (Fig. 2B).

**Figure 2:**
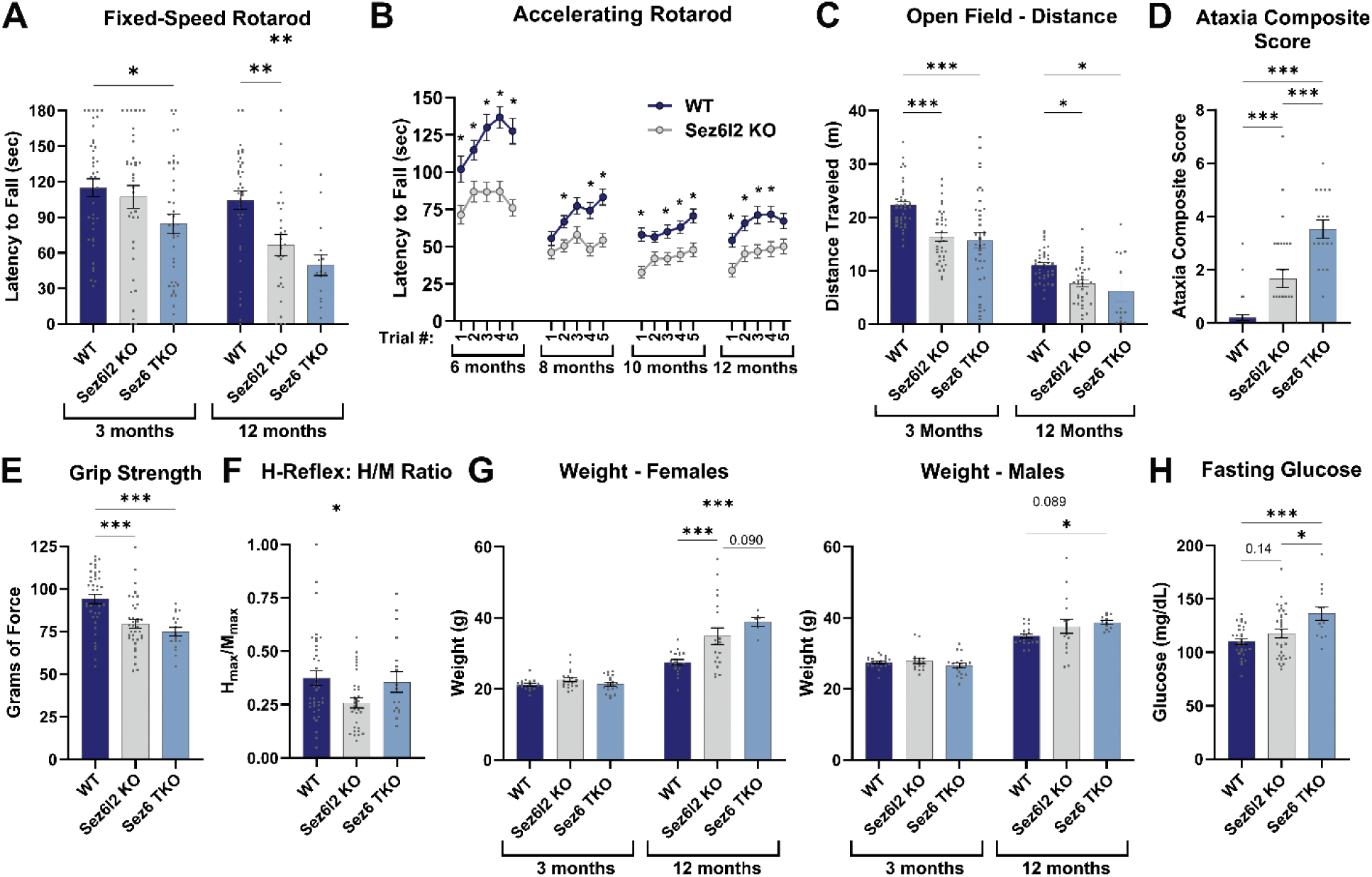
*Sez6l*2 KO and *Sez6* TKO mice show impaired motor function that worsens with age. **A)** Mice were assessed on a fixed speed rotarod for latency to fall. Statistics: 2-Way ANOVA with HS MCTs. 3 months: n=36-40; 12 months: n=15-37. **B)** WT and *Sez6l2* KO mice were evaluated for latency to fall on an accelerating rotarod with 5 trials per timepoint. Statistics: 2-Way ANOVA with HS MCTs; n=38-39. **C)** Quantification of the distance traveled in an open field assay. Statistics: 2-Way ANOVA with HS MCTs. 3 months: n=36-40; 12 months: n=13-38. **D)** Graph of ataxia composite score at 12 months. Statistics: 1-way ANOVA with HS MCTs; n=17-38. **E)** Quantification of forelimb grip strength at 12 months. Statistics: 1-way ANOVA with HS MCTs; n=18-38. **F)** Quantification of H-reflex H/M ratio recorded at 12 months. Statistics: Welch ANOVA with D.T3 MCTs; n=18-37. **G)** Weight. Statistics: 2-way ANOVA with HS MCT; 3 months: n=19-20F, 17-20M; 12 months: n=6-19F, 12-19M. **H)** Fasting glucose measured at 12 months. Statistics: 1-way ANOVA with HS MCT; n=14-34.

At 1 year, motor function was further assessed using the open field test, ataxia composite score, grip strength assessment, and H-reflex assays. *Sez6l2* KO mice showed reduced distance traveled, higher ataxia scores, and weaker grip strength compared to WT mice, with *Sez6* TKO mice exhibiting more severe impairments (Fig. 2C-E). Although we hypothesize many of these deficits are secondary to central nervous system involvement, *Sez6l2* KO mice had a lower H/M wave ratio, the index of excitability in the spinal cord motor neuron pool, suggesting potential peripheral sensory-motor transmission deficits (Fig. 2F). No weight differences were observed at 3 months, but 1-year-old female *Sez6l2* KO and *Sez6* TKO mice were significantly heavier than WT mice, with no differences in males (Fig. 2G). Given the notable weight increase, we measured fasting glucose levels to assess potential metabolic dysfunction that could impact the nervous system. Fasting glucose levels were elevated in *Sez6* TKO mice, with only a minor upward trend observed in *Sez6l2* KO mice (Fig. 2H).

In summary, *Sez6l2* KO mice exhibit intermediate motor impairments relative to WT and *Sez6* TKO mice, highlighting the unique role of *Sez6l2* in maintaining motor function. While all *Sez6* family members contribute to mobility, impairments are exacerbated in TKO mice, indicating limited compensatory ability among family members.

### Low sociability in *Sez6l2* KO mice

Sociability was assessed using a 3-chamber stranger versus object test. While all genotypes appeared to spend more time interacting with the stranger mouse than the object, this difference was statistically significant only for WT and *Sez6* TKO. *Sez6l2* KO showed reduced interaction with the stranger compared to WT and *Sez6* TKO mice (Fig. 3A-C), resulting in a significantly lower social preference index. Additionally, both *Sez6l2* KO and *Sez6* TKO mice exhibited increased immobility during the assay (Fig. 3D).

**Figure 3.**
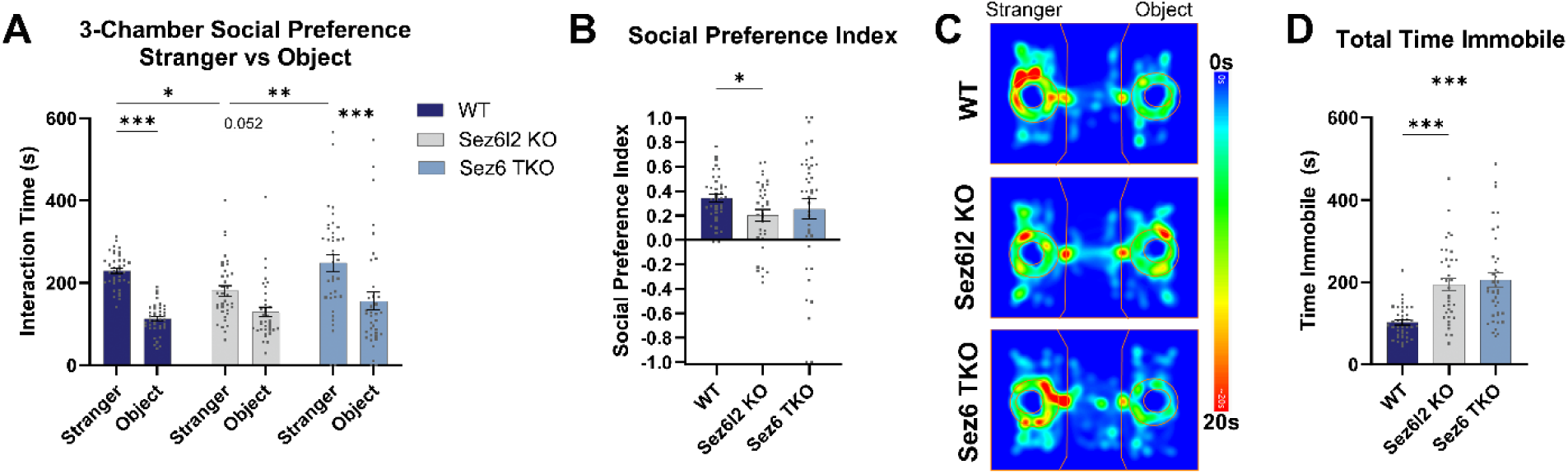
*Sez6l2* KO mice show low sociability in 3-chamber stranger versus object test. **A)** Time mice spent interacting with the stranger mouse versus the object. Statistics: 2-way RM ANOVA with HS MCTs. **B)** Social preference index. **C)** Representative heatmaps from the social preference test. **D)** Total time immobile during the social preference test. Statistics for B and D: Welch ANOVAs with D.T3 MCT. N’s for all graphs: WT n=36-40.

### Reduced Repetitive Behavior in *Sez6l2* KO and *Sez6* TKO Mice

Innate repetitive behaviors were evaluated through marble burying and nesting assays, which reflect both motor and cognitive functions. *Sez6l2* KO mice buried fewer marbles than WT mice, while *Sez6* TKO mice performed worse, taking longer to initiate digging and burying fewer marbles than both WT and *Sez6l2* KO mice (Fig. 4A-C). In the nesting assay, *Sez6l2* KO and *Sez6* TKO mice left nestlets largely intact, producing poor or no nests (Fig. 4D,E). These results suggest that the loss of *Sez6* family genes impairs both repetitive and goal-directed behaviors, emphasizing their role in motor and cognitive functions.

**Figure 4:**
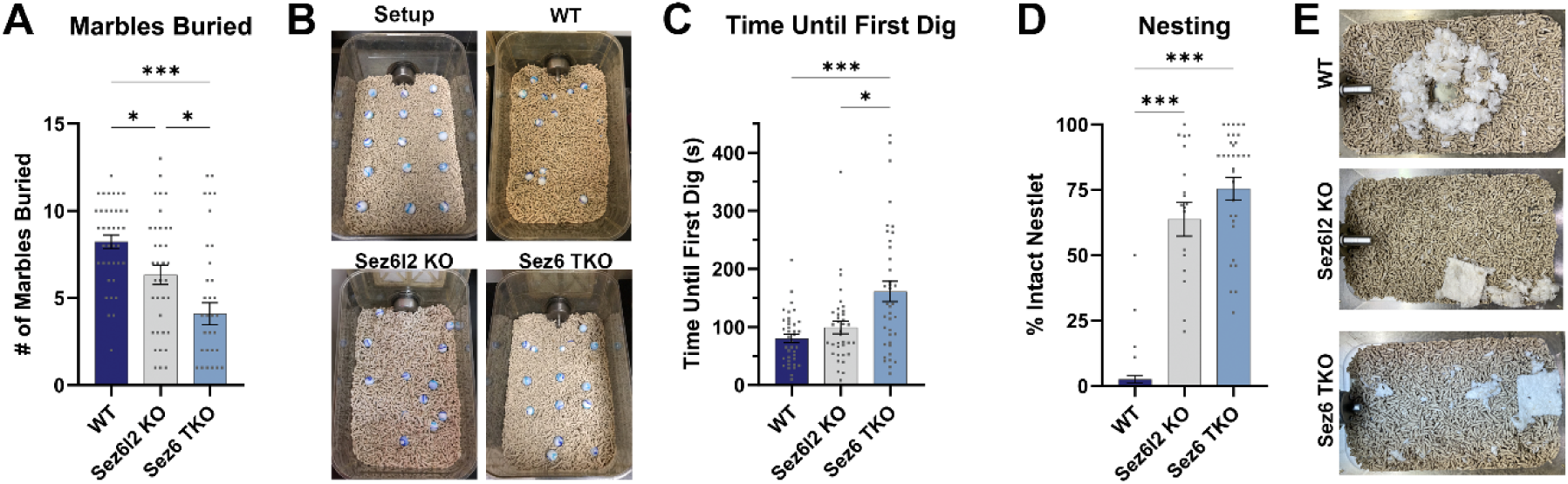
*Sez6l2* KO and *Sez6* TKO demonstrate reduced innate behaviors of repetitive digging (marble burying assay) and nesting. **A)** Number of marbles buried; n=36-40. **B)** Representative images of the marble setup and then marbles buried by WT, *Sez6l2* KO, and *Sez6* TKO mice. **C)** Time until mice started digging; n=36-40. (**D, E)** Nesting assay. **D)** Percent of nestlet that remained intact; n=19-41. **E)** Representative images of nests. Statistics for all graphs: Welch ANOVAs with D.T3 MCT.

### Decreased Auditory Startle Responses and Increased PPI in Male *Sez6l2* KO mice

Auditory startle reflex and prepulse inhibition (PPI), a measure of sensorimotor gating, were assessed. Both male and female *Sez6l2* KO and *Sez6* TKO mice showed significantly reduced startle responses compared to WT mice. However, female *Sez6l2* KO and *Sez6* TKO mice were equally impaired, while male *Sez6l2* KO mice exhibited intermediate startle responses between WT and *Sez6* TKO mice (Fig. 5A). Reduced startle responses have previously been linked to neurodevelopmental, neuropsychiatric, and neurodegenerative diseases, likely reflecting cognitive or motor impairments (Medina et al. 2001; Giakoumaki et al. 2010; Beck 2013; Sichler et al. 2019).

**Figure 5:**
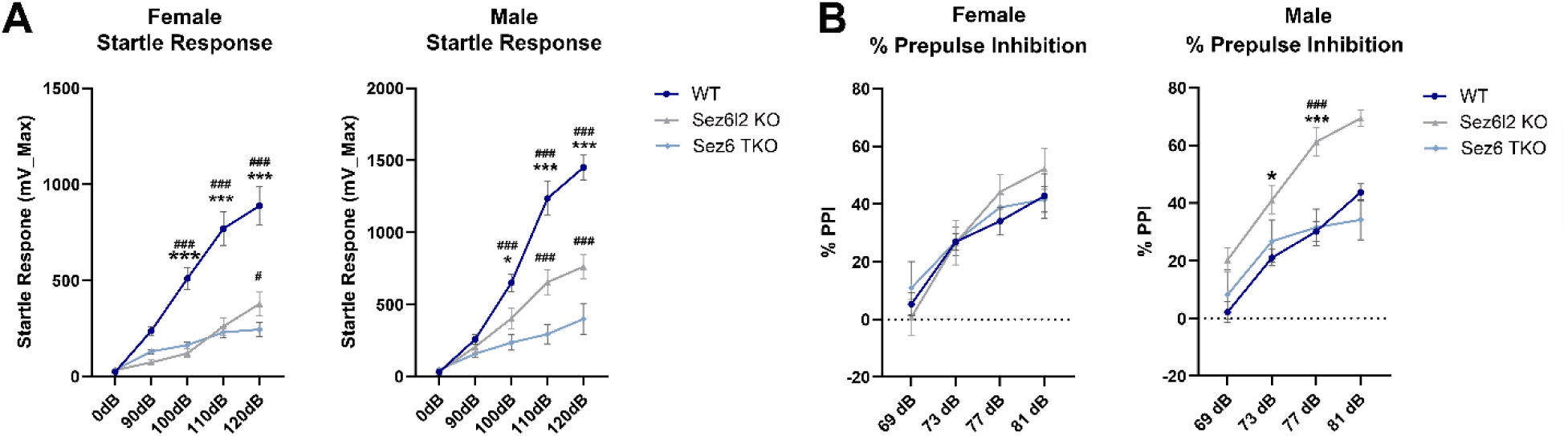
*Sez6l2* KO and *Sez6* TKO have a decreased startle response and only male *Sez6l2* KO mice have increased prepulse inhibition. A) Startle response in response to auditory stimuli of the indicated decibels (dB). B) Percent prepulse inhibition. Statistics for all graphs: 2-way ANOVA with HS MCT; n=36-40. * p<0.05 for comparisons with WT; # p<0.05 for comparisons with *Sez6* TKO.

PPI reflects the nervous system’s ability to filter irrelevant stimuli by reducing the startle response to a strong stimulus when preceded by a weaker prepulse. All genotypes showed the expected reduced startle responses (increased % PPI) when preceded by prepulse stimuli. However, male *Sez6l2* KO mice exhibited significantly higher PPI compared to WT and *Sez6* TKO mice (Figure 5B). This suggests a male-specific alteration in brain function linked to enhanced filtering of irrelevant stimuli or a compensatory mechanism for their reduced startle response.

### Heightened Fear Memory in *Sez6l2* KO Mice and Auditory Cued Fear Deficits in *Sez6* TKO Mice

Contextual and cued fear conditioning assays were used to assess memory impairments. Male *Sez6l2* KO mice froze significantly more in the conditioned context compared to WT or *Sez6* TKO males, while female *Sez6l2* KO and *Sez6* TKO showed a non-significant trend of increased freezing (Fig. 6A). In a novel context with the conditioned tone, male *Sez6l2* KO mice again froze more than WT and *Sez6* TKO males (Fig. 6B-C). Surprisingly, male and female *Sez6* TKO mice froze significantly less in response to the tone than WT and *Sez6l2* KO mice. Minimal freezing was observed across all genotypes in the novel context before the tone, indicating recognition of the altered environment (Fig. 6C). These results suggest male *Sez6l2* KO mice exhibit heightened fear memory in both contextual and cued paradigms, while *Sez6* TKO mice show deficits in the auditory cued fear test.

**Figure 6:**
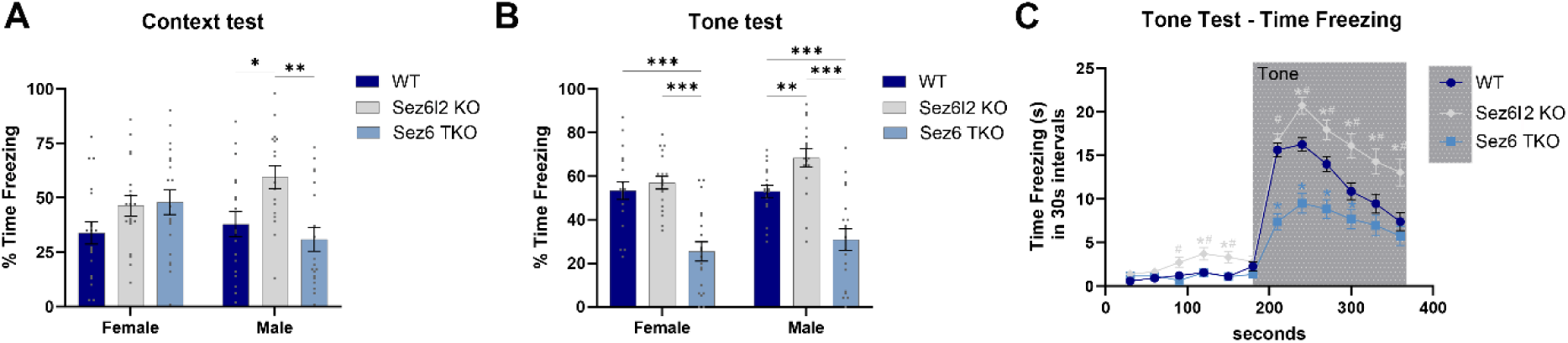
*Sez6l2* KO and *Sez6* TKO mice show sex-dependent memory deficits in a contextual fear assay. **A)** % time freezing in the conditioned context test. **B)** % time freezing in the novel context after the tone. **C)** Time freezing in 30s intervals in the combined novel context/tone test. C includes both male and female mice. Statistics for all graphs: 2-way ANOVA with HS MCT; n=36-38 (19F, 17-19M). In C, * p<0.05 for comparisons with WT. # p<0.05 for comparisons with *Sez6* TKO.

### Auditory Impairment in *Sez6* TKO Mice

To determine if auditory deficits contributed to behavioral differences, hearing thresholds were assessed using auditory brainstem response (ABR) and distortion product otoacoustic emissions (DPOAE) assays. *Sez6* TKO mice exhibited significantly higher ABR thresholds across all tested frequencies, indicating impaired auditory sensitivity (Fig. 7A). However, their DPOAE measurements were comparable to WT, suggesting normal cochlear function (Fig. 7B). *Sez6l2* KO mice had ABR thresholds identical to WT mice, indicating normal auditory sensitivity. These findings confirm auditory impairment in *Sez6* TKO mice but provide no evidence that auditory sensitivity is a confounding factor in *Sez6l2* KO behavioral phenotypes.

**Figure 7:**
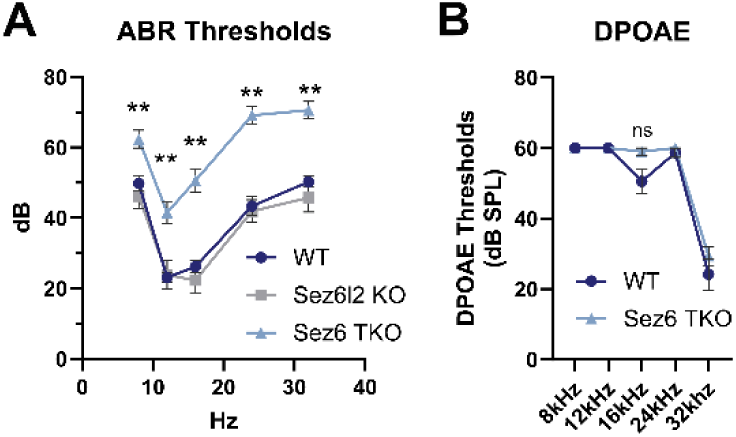
*Sez6* TKO mice have significant hearing impairments. **A)** Mean auditory brainstem response (ABR) thresholds at the indicated frequencies and decibels. **B)** Mean distortion product otoacoustic emissions (DPOAE) thresholds. Statistics: 2-Way ANOVA with HS MCTs; n=13-21.

### Synaptic Changes in *Sez6l2* KO and *Sez6* TKO mice

To determine whether behavioral abnormalities in *Sez6l2* KO and *Sez6* TKO mice are linked to synaptic changes, we analyzed dendritic spine density and spine length in the somatosensory cortex at 3 months using Golgi staining. While overall spine density was unchanged in both genotypes, spine length showed significant alterations (Fig. 8). Using a modified Rapid Golgi Analysis Method (Risher et al. 2014), spines were classified as long (>2.4 μm), medium (1–2.4 μm), or short (<1 μm). Both *Sez6l2* KO and *Sez6* TKO mice exhibited fewer long spines, with *Sez6* TKO mice also showing fewer medium spines and an increase in short spines (Fig 8B). Cumulative spine length analysis revealed a shift toward shorter spines in both genotypes, with *Sez6l2* KO spine lengths intermediate between WT and *Sez6* TKO (Fig. 8C).

**Figure 8:**
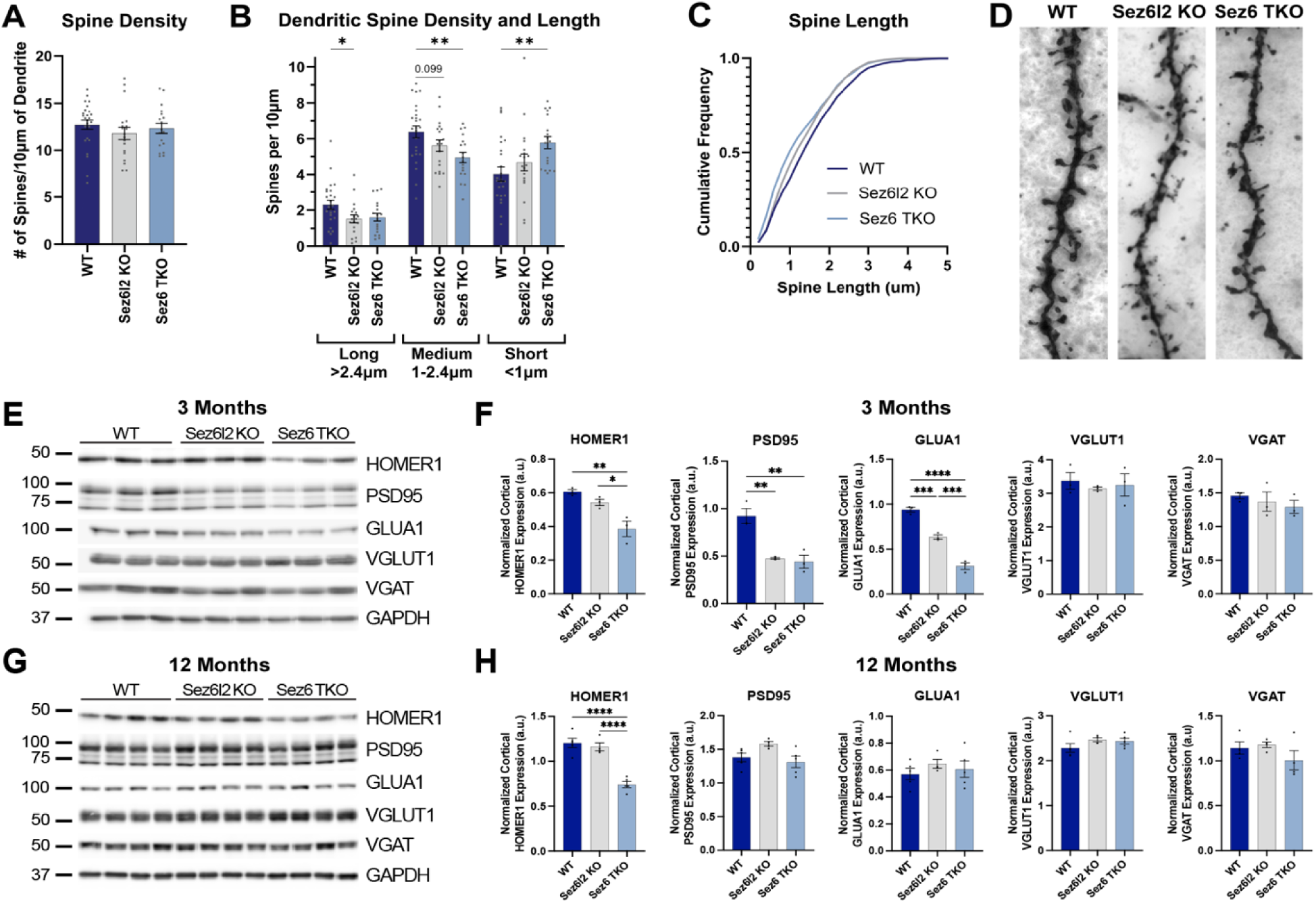
*Sez6l2* KO and *Sez6* TKO mice exhibit altered cortical synaptic architecture. **A)** Quantification of spine density. **B)** Quantification of spine density binned by length. Statistics for A and B: 1 or 2-way ANOVAs with HS MCT; n=18-25 dendrites per group, with 2-4 dendrites quantified per mouse. **C)** Cumulative Frequency of spine length; n=1395-1895 spines. **D)** Representative Golgi-stained dendrites. Images are 10x40 μm. (**E-H)** Synaptic protein expression was measured by western blot in cortical lysates from (E, F) young adult (3 months) and (G,H) aged (12 months) WT, *Sez6l2* KO, and *Sez6* TKO mice and normalized to GAPDH. n=3-4 mice per genotype. Statistics 1-way ANOVA with Tukey’s MCT.

To assess synaptic protein changes, cortical lysates were analyzed by western blot. At 3 months of age, *Sez6l2* KO and *Sez6* TKO mice showed reduced expression of excitatory postsynaptic markers PSD95 and GLUA1, with *Sez6* TKO mice also exhibiting lower HOMER1 levels (Fig. 8E,F). Presynaptic markers VGLUT1 (excitatory) and VGAT (inhibitory) remained unchanged. By 12 months, PSD95 and GLUA1 levels returned to WT levels, but HOMER1 remained reduced in *Sez6* TKO mice (Fig. 8G,H).

The observed reduction in dendritic spine length and selective decreases in synaptic proteins provide a potential neurological correlate for the motor, cognitive, and sensory impairments seen in *Sez6l2* KO and *Sez6* TKO mice.

### Genetic loss of complement component C3 partially rescues spine architecture and select gait deficits in *Sez6* TKO mice but does not broadly improve behavioral outcomes

SEZ6L2 has been reported to inhibit complement deposition on cell surfaces (Qiu et al., 2021). Given that complement C3b mediates synaptic pruning by microglia in development (Stevens et al., 2007), loss of SEZ6 family proteins may lead to aberrant complement activation and synaptic refinement, contributing to reduced dendritic spine size and synaptic protein expression in *Sez6* TKO mice. To test this, *Sez6* TKO mice were crossed with *C3* KO mice to generate *Sez6* TKO/*C3* KO animals, and spine morphology and behavior were compared across genotypes.

Loss of C3 significantly rescued spine length deficits in the somatosensory cortex of *Sez6* TKO mice, with *Sez6* TKO/*C3* KO and *C3* KO mice resembling WT (Figure 9 A-D). Gait analysis showed modest improvement: *Sez6* TKO/*C3* KO mice exhibited a trend toward reduced hindlimb base widening and improved step sequence regularity compared to *Sez6* TKO mice, along with a shorter run duration (Figure 9 E-G). These effects were independent of body weight, although *Sez6* TKO mice gained more weight by 1 year, a phenotype not observed in *Sez6* TKO/*C3* KO mice (Figure 9 H).

**Figure 9:**
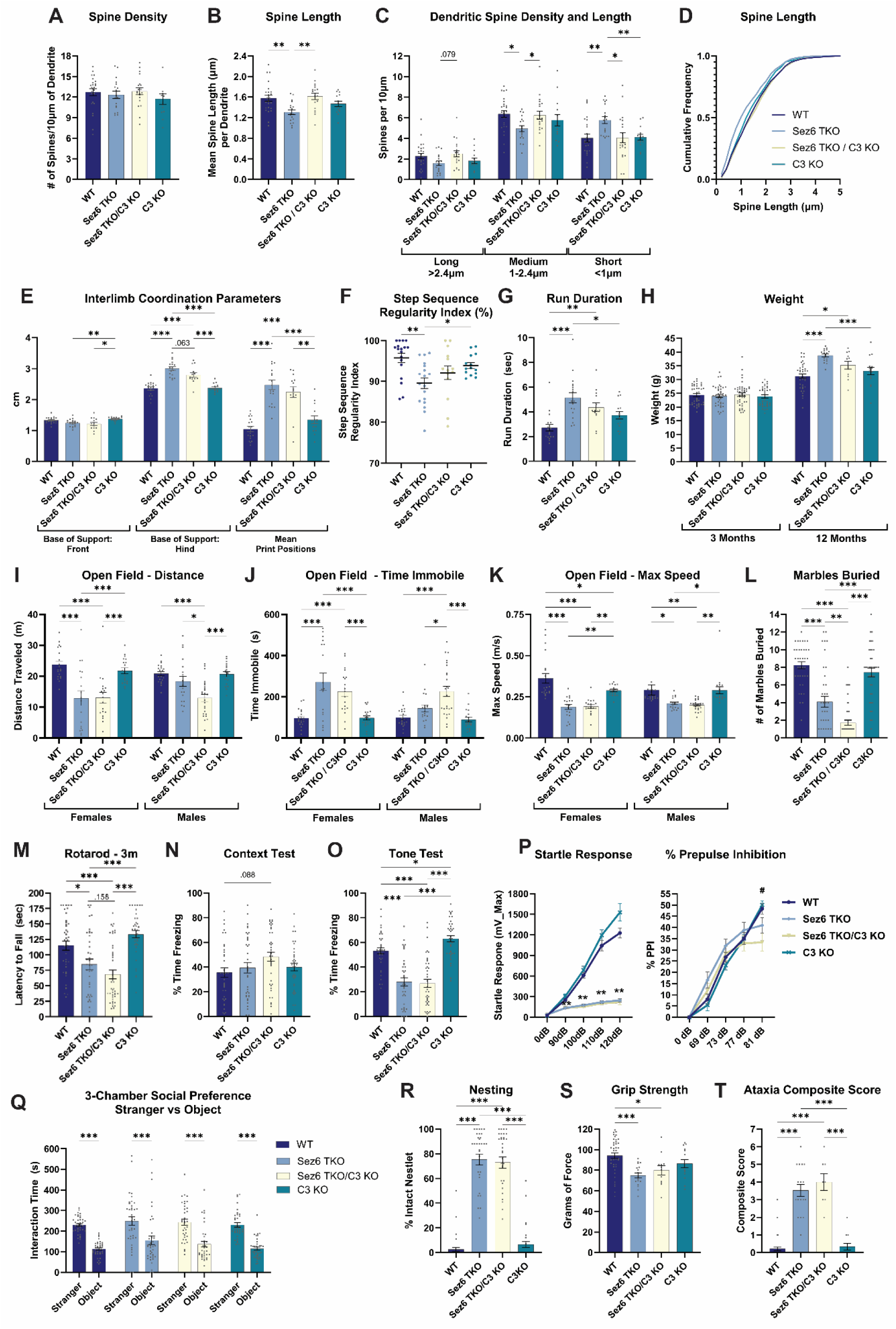
Genetic loss of complement component C3 protects spine architecture and some gait impairments in *Sez6* TKO mice; but does not improve other behavior outcomes. All data is from mice at 3-4 months of age unless stated otherwise. **A-C)** Quantification of mean spine density (A) and Length (B) per dendrite. In C the spine density is binned by length. Statistics for A-C: 1 or 2-way ANOVAs with HS MCT; n=dendrites per group (WT = 25, *Sez6* TKO = 18, *Sez6* TKO/*C3* KO = 20, *C3* KO = 14) with 2-4 dendrites quantified per mouse. **D)** Cumulative Frequency of spine length; n=979-1895 spines per group. **E-G)** Gait analysis metrics: **E)** interlimb coordination parameters, **F)** step sequence regularity index, **G)** run duration. Statistics for E-G: n=14-18 mice per group; E: 2-way repeated-measure ANOVAs with HS MCTs; F: Welch ANOVA with D.T3 MCT; for G: 1-way ANOVA with HS MCTs. **H)** Average weight at 3 and 12 months. Statistics: 2-Way ANOVA with HS MCT; 3 months: 38-45; 12 months n=12-38. **I-K)** Quantification of the distance traveled (I), time immobile (J), and max speed (K) in an open field assay at 3 months. Statistics: 2-Way ANOVA with HS MCTs: n=19-20F, 19-25M. **L)** Number of marbles buried. Statistics: Welch ANOVA with D.T3 MCT; n=38-47. **M)** Latency to fall on a fixed-speed rotarod at 3 months. Statistics: 1-Way ANOVA with HS MCTs; n=36-45. **N-O)** Percent time freezing in the context test (N) or tone test (O) 1 day following contextual fear conditioning. Statistics: Statistics: 1-Way ANOVA with HS MCTs; n=37-41. **P)** Startle response in response to auditory stimuli of the indicated decibels (dB) and percent prepulse inhibition. Statistics: 2-way ANOVAs with HS MCT; n=38-44. **Q)** Time mice spent interacting with the stranger mouse versus the object in a 3-chamber sociability test. Statistics: 2-way RM ANOVA with HS MCTs. **R)** Percent of nestlet that remained intact in a 24-hour nesting assay; Statistics: Welch ANOVA with D.T3 MCT; n=32-41. **S)** Quantification of forelimb grip strength at 12 months. Statistics: 1-way ANOVA with HS MCTs; n=12-38. **T)** Graph of ataxia composite score at 12 months; Statistics: 1-way ANOVA with HS MCTs; n=8-38. For all graphs in Figure 9: the WT and *Sez6* TKO data are the same as shown in the previous figures.

Despite these structural and partial motor improvements, behavioral outcomes were not broadly rescued. *Sez6* TKO/*C3* KO mice showed worsened performance in some assays, including reduced locomotion in the open field (males), decreased marble burying, and trends toward impaired rotarod performance and increased contextual freezing (Figure 9 I-O). Other measures, including behavioral responses to sound, prepulse inhibition, sociability, nesting, grip strength, and composite ataxia scores, were unchanged between *Sez6* TKO and *Sez6* TKO/*C3* KO mice (Figure 9 P-T). Together, these findings indicate that complement C3 contributes to synaptic and select motor deficits in *Sez6* TKO mice but is not a primary driver of broader behavioral impairments.

## DISCUSSION

### SEZ6 Proteins and Sensorimotor Impairments

We observed significant motor impairments in *Sez6l2* KO mice in a variety of motor tasks, including rotarod, gait analysis, open field, and grip strength assays. Similarly, *Sez6* TKO mice also displayed these motor deficits, often with a more severe impairment than that of the single KO model. These findings are consistent with previous observations in *Sez6* TKO mice (Miyazaki et al. 2006; Nash et al. 2020), but also provide the first evidence of motor impairments in the single knockout model of *Sez6l2*. Abnormal motor function is a common phenotype present in neurodevelopmental disorders such as ASD. Children with ASD often display gross or fine motor skill impairments, as well as abnormal locomotion and coordination (Vernazza-Martin et al. 2005; Provost et al. 2007). In mouse models of ASD, mice present with irregular gait, coordination dysfunction, and motor learning deficits (Piochon et al. 2014; Cording and Bateup 2023). Moreover, the 16p11.2 deletion (16p11.2 DEL/+) model of ASD, which includes a hemizygous deletion of *Sez6l2,* presents with tremors, motor delays, and gait abnormalities (Portmann et al. 2014). These motor phenotypes may reflect a dysfunction in proprioception and abnormal sensorimotor coordination linked to improper neurodevelopment. Our findings reinforce the critical role of SEZ6 family proteins in motor function and emphasize SEZ6L2 as a key contributor to the motor deficits observed in *Sez6* TKO mice, and likely in 16p11.2 DEL/+ mice as well.

Hearing impairments were found in *Sez6* TKO mice, as evidenced by elevated ABR thresholds despite normal cochlear function. In contrast, *Sez6l2* KO mice did not exhibit this same decrease in auditory sensitivity at the tested frequencies. Thus, the auditory deficit must be due to the loss of either *Sez6* or *Sez6l* alone or require the loss of multiple family members. Thus far, only *Sez6* has been identified as a candidate for recessive hearing impairment in humans (Bharadwaj et al. 2022), suggesting that it might be the critical family member. This auditory deficit is particularly important for interpreting the behavioral observations of *Sez6* TKO and *Sez6* TKO/C3 KO mice, especially in assays involving auditory stimuli such as PPI and tone-cued fear conditioning assays.

This study also provides evidence of abnormal sensorimotor responses in *Sez6l2* KO mice, as they exhibited a reduced acoustic startle response relative to WT. This reduction may partially be a consequence of the observed motor impairments, or it may indicate changes in sensorimotor gating and integration. In male *Sez6l2* KO mice, this altered sensory processing is further reflected by the elevated PPI response. PPI is an established behavioral measurement for sensorimotor gating and is often disrupted in neurological disorders. The PPI response is most studied as a “biomarker” for schizophrenia, where the attenuation of prepulse inhibition is directly correlated with the severity of positive schizophrenia symptoms (Weike et al. 2000; Mena et al. 2016). On the other hand, PPI is often elevated in ASD patients, indicating hypersensitivity to sensory stimuli (Madsen et al. 2014). Certain mouse models of ASD, such as the *Cntnap* KO mouse, also exhibit increased PPI (Brunner et al. 2015). Reports of PPI abnormalities in 16p11.2 DEL/+ mice, however, are mixed, with some studies showing attenuated PPI, others finding only sex-specific deficits, and others observing intact PPI responses (Brunner et al. 2015; Lynch et al. 2020; Openshaw et al. 2023). Despite the variation in observed PPI phenotypes across disorders and mouse models, it is likely that PPI alterations indicate a disruption in sensorimotor gating and processing. This behavior, therefore, provides further evidence that SEZ6L2 is important for normal sensory integration.

Some behavioral phenotypes in our *Sez6l2* KO and *Sez6* TKO mice were sex dependent. Male *Sez6l2* KO mice showed heightened fear learning responses in both the contextual and tone-cued assessments, reflecting increased fear memory and an anxiety-like phenotype. Enhanced PPI responses were also limited to males. Similar sex-related phenotypic differences are observed in humans with neurodevelopmental disorders (Werling and Geschwind 2013).

### SEZ6 Proteins in Social and Repetitive Behaviors

Abnormal social behaviors are also a common hallmark of neurodevelopmental and neuropsychiatric disorders in both humans and mice (Baranek 1999; Kazdoba et al. 2016; Bishop-Fitzpatrick et al. 2017; Benedetti et al. 2022). In this study, we demonstrate that *Sez6l2* KO mice have a reduced social preference index compared to WT mice. The 16p11.2 DEL/+ mouse model has also been shown to exhibit abnormal social behaviors across various sociality assays and environmental observations (Arbogast et al. 2016; Rusu et al. 2023). However, other studies have reported that the genetic background of the 16p11.2 DEL/+ model significantly modifies this phenotype, with some models displaying no observable social deficits (Horev et al. 2011; Benedetti et al. 2022). Our findings suggest that the heterozygous loss of *Sez6l2* likely contributes to the social impairments observed in 16p11.2 DEL/+ mice; however, it may not be sufficient to drive these behaviors on its own. Surprisingly, *Sez6* TKO mice did not display the social deficits observed in *Sez6l2* KO mice, though the *Sez6* TKO group showed more variability in their results. For example, several *Sez6* TKO mice exhibited a strong negative social preference or only visited one end of the chamber, suggesting they may have failed to engage in the social test altogether. Perhaps, the more pronounced motor deficits and anxiety-like behavior in the *Sez6* TKO mice also partially confounded the ability to accurately measure their social behavior.

We found that *Sez6l2* KO and *Sez6* TKO mice exhibited a reduction in marble burying and nestlet shredding behaviors. Marble burying and nestlet shredding represent goal-directed behaviors, which can be modulated by glutamatergic signaling. Targeting glutamatergic receptors has repeatedly been found to decrease these repetitive innate behaviors (Iijima et al. 2010; de Brouwer et al. 2019; Ma et al. 2023). Based on our findings of decreased GLUA1 expression in both *Sez6l2* KO and *Sez6* TKO mice at the age of testing, these changes in repetitive goal-directed behaviors may be explained by a reduction in glutamatergic activity. It is also possible that this finding is a result of the motor deficits in our mice, rather than a direct impact on repetitive behaviors. The deficits in measures such as interlimb coordination, locomotion, and maximum speed observed in our *Sez6l2* KO and *Sez6* TKO mice indicate a moderate ataxic phenotype that may lead to fewer or less efficient coordinated movements that would be required for the marble burying and nestlet shredding tasks. Together, our findings indicate that there may be a motor component, as well as a reduction in the drive to initiate repetitive movements, that underlie the deficits in these behavioral tasks.

This decrease in repetitive innate behaviors was initially unexpected, as neurodevelopmental disorder phenotypes are often associated with increased repetitive activity in both human studies and mouse models (Baranek 1999; Whitehouse and Lewis 2015; Gandhi and Lee 2020). However, mouse models of certain neuropsychiatric disorders, such as schizophrenia, have been reported to exhibit reduced repetitive behaviors (Ma et al. 2023). Deficits in nest-building behaviors have also been observed early in the phenotypic characterization of mouse models of Alzheimer’s disease (AD) (Filali and Lalonde 2009). Interestingly, a recent transcriptomic analysis study across multiple AD datasets identified the downregulation of *Sez6l2* to be a significant risk factor for the development of AD (Nguyen et al. 2024). Furthermore, genomic analyses have shown genes related to synaptic processes to be key risk factors for both neurodevelopmental disorders and neurodegenerative diseases (Selkoe 2002; De Rubeis et al. 2014; Lepeta et al. 2016; Wilfert et al. 2017), indicating a potential overlap in risk genes. Although some motor dysfunction was already present in young *Sez6l2* KO adult mice, our data show that the motor impairments worsen with age. These findings further implicate SEZ6 family proteins in the pathophysiology of neurodegenerative disorders characterized by age-dependent motor decline, such as amyotrophic lateral sclerosis (ALS) or Parkinson’s disease. Based on these findings, SEZ6 proteins likely play an important role in the pathology of disorders of both development and degeneration.

### SEZ6 Family Proteins and Synaptic Development

Based on previous studies showing SEZ6 proteins to be important in synaptic architecture and function (Miyazaki et al. 2006; Gunnersen et al. 2007; Gunnersen et al. 2009; Zhu et al. 2018; Nash et al. 2020), we hypothesized that the behavioral differences identified in the *Sez6l2* KO and *Sez6* TKO mouse models have a neurological origin based on alterations in synaptic connectivity. We identified changes in both dendritic spine morphology and synaptic protein expression as structural correlates that may underlie these observed behaviors. Analysis of dendritic spines revealed a decrease in spine length in both *Sez6l2* KO and *Sez6* TKO mice, consistent with previous findings by Nash and colleagues that loss of SEZ6 proteins leads to increased frequency of short, stubby spines (Nash et al. 2020).

We also observed selective reductions in synaptic proteins in the knockout groups. At 3 months, both knockout strains showed decreased expression of post-synaptic markers PSD95 and GLUA1. SEZ6L2 is a putative direct binding partner GLUA1 and may regulate AMPA receptor trafficking and/or stabilization (Yaguchi et al. 2017), providing a potential explanation for reduced GLUA1 levels in the absence of SEZ6L2. Decreased AMPAR signaling may in turn contribute to reduced spine size, as AMPAR-dependent activity is required for spine growth during long-term potentiation (Matsuzaki et al. 2004). Similarly, reduced PSD95 and HOMER1 levels in our knockout mice may further impair synaptic maturation and stability as the knockdown of PSD95 or HOMER1 prevents LTP-induced spine growth and maturation (Ehrlich et al. 2007; Gerstein et al. 2012); (Yoon et al. 2021).

HOMER1 expression was persistently reduced in *Sez6* TKO mice but not *Sez6l2* KO mice, suggesting more sustained synaptic scaffolding deficits in the TKO model. In contrast, PSD95 and GLUA1 levels appeared to normalize by 12 months, possibly reflecting compensatory or experience-dependent changes. However, given the continued behavioral decline with age, these molecular changes are unlikely to indicate true functional recovery.

Another explanation for these discrepancies may be due to environmental differences between the two cohorts of mice. The cortical lysates generated from 12-month-old mice were collected from the cohort that underwent the battery of behavioral assays, while the lysates from 3-month-old mice, as well as the brains used for the Golgi stain, were collected from behaviorally naïve mice. Behavioral assessments, especially repeated motor tasks, can serve as an enriched environment, which has been shown to increase dendritic spine density (Jung and Herms 2014). This increase can lead to elevated expression of synaptic proteins PSD95 and GLUA1 (Ehrlich et al. 2007; Shinohara 2012), which might appear as a “recovery” to wildtype levels. Finally, another interpretation of these results may be that the shifting levels of PSD95 and GLUA1 are indicative of a decrease in dendritic spine stability. Dendritic spine destabilization, characterized by prolonged spine turnover and reduced mature spine morphologies, is associated with many neurodevelopmental and neuropsychiatric disorders (Cruz-Martin et al. 2010; De Rubeis et al. 2013; Jiang et al. 2013). If the loss of SEZ6 proteins similarly leads to destabilization of dendritic spines, this may contribute to the fluctuations in synaptic protein expression observed here.

One proposed mechanism linking SEZ6 proteins to synaptic structure is their role in regulating complement activity. SEZ6L2 and SEZ6L have been shown to inhibit complement opsonization of cell surfaces (Qiu et al. 2021). As complement C3 mediates synaptic pruning during development (Stevens et al. 2007), loss of all SEZ6 proteins may lead to excessive complement-mediated synapse elimination. Consistent with this model, genetic deletion of C3 in *Sez6* TKO mice rescued dendritic spine length and partially improved select gait parameters. However, broader behavioral deficits were not improved and were in some cases modestly exacerbated. Our synapse analyses were limited to cortical regions, with spine morphology measured in the somatosensory cortex and synaptic protein expression in cortical lysates. These measures may not reflect changes in other behaviorally relevant circuits. Together, these findings indicate that complement dependent synaptic remodeling contributes to structural and select motor phenotypes but does not account for the broader behavioral impairments in *Sez6* mutant mice, which likely arise from additional mechanisms including disrupted glutamatergic signaling, altered synaptic scaffolding, and spine instability.

In summary, our results identify SEZ6L2 as an essential regulator of synaptic integrity and behavior, with loss of function leading to progressive neurological deficits. The overlap between synaptic, motor, and behavioral abnormalities observed here and those seen in neurodevelopmental and neurodegenerative disorders highlights SEZ6 family proteins as important contributors to disease-relevant pathways. These findings provide a foundation for future studies aimed at targeting SEZ6-mediated mechanisms to restore synaptic function.

## MATERIALS AND METHODS

### Mice

Animal care and use were carried out in compliance with the US National Council’s Guide for the Care and Use of Laboratory Animals and the US Public Health Service’s Policy on Humane Care and Use of Laboratory Animals. Protocols were approved by the University Committee on Animal Resources at the University of Rochester.

The *Sez6l2* KO (*Sez6l2*^tm1Hta^, MGI:3665419) and *Sez6* TKO (*Sez6*^tm1Hta^, *Sez6l*^tm1Hta^, *Sez6l2*^tm1Hta^, MGI:3688101) mouse lines were generously provided by Dr. Jenny Gunnersen and Dr. Hiroshi Takeshima (Miyazaki et al. 2006; Nash et al. 2020). Knockout of SEZ6 proteins was verified by western blot (Supplemental Fig. 1). *Sez6l2* mice were backcrossed to the C57BL/6 strain, while *Sez6* TKO mice were maintained on a mixed 129/C57BL/6 background. C57BL/6 and *C3* KO (B6.129S4-*C3^tm1Crr^*/J, MGI:J:30152; Wessels et al. (1995)) mice were obtained from JAX (Strain #000664 and #029661, respectively). *Sez6* TKO mice were crossed with *C3* KO mice to generate the *Sez6* TKO/*C3* KO line. Mice were bred at the University of Rochester and were placed on a reverse 12-hour (hr) light/dark cycle. All behavior experiments were conducted during the dark cycle under low light unless otherwise specified.

### Behavior Testing Schedule

The full behavioral cohort (WT n=40 [20 female (F), 20 male(M)], *Sez6l2* KO n=36 [19F, 17M], *Sez6* TKO n=39 [20F, 19M], *Sez6* TKO/*C3* KO n=47 [21F, 26M], and *C3* KO n=39 [19F, 20M]) was divided into 7 age-based groups with both sexes and mixed genotypes. Weekly behavior assays were conducted by experimenters blinded to genotype in this order (approximate ages in weeks): Nesting (8), move to reverse light cycle, open field (11), marble burying (12), rotarod (13), prepulse inhibition (14), sociability (15), contextual fear conditioning (16), and gait analysis (17-19). Mice were aged up to 1 year, with accelerating rotarod at 6, 8, 10, and 12 months. At 12 months, tests included grip strength, fixed-speed rotarod, open field, ataxia, glucose levels, and H-reflexes.

### Nesting Assay

Mice were singly housed for 24 hrs with a pre-weighed nestlet. Intact nestlet pieces (≥0.1 g) were collected, weighed, and recorded as a percentage of the original weight.

### Open Field Test

Mice were placed in a chamber for 10 minutes (min) and movements were recorded and analyzed with ANY-maze software (Stoelting Co.).

### Marble Burying

Mice were placed individually in a cage with ∼5 cm of bedding and 16 marbles for 30 min. under red lighting. Digging latency and the # of marbles buried ≥50% were recorded.

### Rotarod Test

Mice underwent three 3-minute training sessions on a rotarod (Columbus Instruments, Economex) at 7, 10, and 12 rotations per min (RPM). Mice that fell off were placed back on the rod. A final 1-min training at 12 RPM occurred the next day, followed by three 3-min test trials at 12 RPM, recording latency to fall. For accelerating rotarod assays (Panlab, Harvard Apparatus, 76-0770), mice had 1–3 re-training sessions at 5 RPM for 1 min, followed by five trials in accelerating mode (4–40 RPM over 5 min.).

### Acoustic Startle Response and Prepulse Inhibition (PPI)

Mice were acclimated to the chamber with 65 dB background noise for 15 min over two days. Testing consisted of 3 blocks: 1) 7 trials of 120 dB startle pulses (40 ms) with 10–20 s inter-trial intervals; 2) 25 trials with prepulses (0, 69, 73, 77, or 81 dB; 20 ms) followed by a 100 ms interval and a 120 dB startle pulse; 3) 25 trials of startle pulses at varying volumes (0, 90, 100, 110, or 120 dB; 40 ms). Startle responses (SR) were recorded as maximum force (mV_max) and PPI was calculated as the (mean SR_pulse_ – mean SR _prepulse +pulse_) / mean SR_pulse._

### Three-Chamber Sociability Test

The subject mouse was habituated to the 3-chamber apparatus for 10 min. Then, a stranger mouse (same sex; habituated) was placed under a wire cage in one chamber, and a simple object under a wire cage in the opposite chamber. The subject mouse explored the apparatus for 10 minutes, with time spent in each chamber and sniffing each cage recorded.

### Contextual Fear Conditioning

On day 1, mice were conditioned in a chamber with an electrified grid floor (Actimetrics). After 3 min of no stimulation, they received three rounds of a 15 s white noise followed by a 2 s foot shock (0.19 mA), separated by 30 s. After 1 additional min, mice were returned to their home cage. Approximately 24 hours later, mice underwent a 5-min context test in the same chamber without shocks. A novel context test followed ∼4 hrs later in a round cage with smooth plastic walls, clean bedding, ethanol odor, and red lighting to create a distinct environment. After 3 mins of exploration, a continuous white noise tone was presented for 3 mins. Freezing behavior (immobility ≥0.5 seconds) was analyzed using ANY-maze software (Stoelting Co.). Freezing was quantified in 30-second intervals and as percentages of total test time during specific intervals: 30–120 seconds in the conditioned context, 30–120 seconds in the novel context, and 0–60 seconds during tone presentation.

### Gait Analysis

Gait was assessed using the Noldus CatWalk XT system. Mice crossed a glass platform until 3 compliant runs (uninterrupted motion with ≥3 step cycles; 0.5–10 seconds) were obtained. Runs were conducted in the dark and manually reviewed, with data averaged across the 3 runs per mouse.

### Grip Strength Test

Mice grasped a grip strength meter with their forelimbs and were pulled backward by the tail until releasing the bar. Peak force was recorded over 3 trials per mouse.

### Ataxia Composite Score

Mice were scored for clasping, kyphosis, gait alterations, and performance on a short ledge test as previously described (Guyenet et al. 2010).

### H-Reflex Electrophysiology Measurements

Mice under 2-4% isoflurane anesthesia, had electrodes inserted into the gastrocnemius and proximal to the popliteal fossa, with a ground electrode in the contralateral hindlimb. Electrical stimuli (up to 100 mA, 1 ms) were applied to depolarize tibial nerve fibers, and compound muscle action potentials were recorded. M-wave (early response evoked by efferent motor axons) and H waves (late response evoked by efferent motor axons due to activation by sensory afferents) amplitudes and latencies were recorded and H/M wave amplitude ratios were calculated.

### Auditory Brainstem Response (ABR) and Distortion Product Otoacoustic Emissions (DPOAE)

ABR and DPOAE were measured under ketamine and xylazine (100 and 3 mg/kg, respectively) anesthesia using a Smart EP Universal Smart Box (Intelligent Hearing Systems) with Tucker Davis Systems speakers. Subcutaneous electrodes were placed ventral to the tested ear, at the crown, and at the tail (ground). A 10B+ probe delivered 5 ms tone pips (10–95 dB, 8–32 kHz) and ABR responses were averaged over 512 sweeps. Then anterior electrodes were removed for DPOAE testing. Otoacoustic emissions were measured for paired tones (f1/f2 = 1.2, f1 10 dB above f2) across 32 sweeps in 5 dB steps (20–65 dB, 8–32 kHz). DPOAE thresholds were calculated by interpolating the f2 level that produced a 3 dB response. The ABR and DPOAE measurements were taken on a separate cohort of mice from the one used for behavior analysis.

### Golgi Stain

Behavior naïve mice (17-18 week old) were anesthetized with ketamine and xylazine (100 and 10 mg/kg, respectively) and perfused with PBS. Half-brains were collected and stained using the FD Rapid Golgi Stain Kit (FD NeuroTechnologies Inc., PK401A). 100 μm sections were imaged on a Nikon A1R HD confocal microscope (60x) to visualize layer IV/V neuronal secondary dendrites projecting into layer II/III of the somatosensory cortex. Dendritic spines were analyzed using Imaris software with the filament tracer tool and manual adjustments.

### Western Blots

Cortical halves from 3.5- or 12-month-old mice were processed for western blotting as previously described (Reyes-Sepulveda et al. 2025). Primary antibodies used included HOMER1 (Synaptic Systems), PSD-95 (DSHB, K28/43), GLUA1 (MilliporeSigma, MAB2263), VGLUT1 (MilliporeSigma, AB5905), VGAT (Synaptic Systems, 131004), and GAPDH (MilliporeSigma, CB1001) with HRP secondary antibodies from Azure Biosystems. Band densities were quantified using FIJI and normalized to the GAPDH loading control.

### Fasting Glucose Test

After a 5-hour fast (starting 5 hours after the light-to-dark switch), tail vein blood glucose levels were measured using a Contour Next One meter and test strips.

### Statistical Analysis

Statistical analyses were performed using GraphPad Prism (La Jolla California USA). Behavior data were analyzed with 2-way ANOVAs to assess sex differences and genotype-sex interactions. Data is displayed separately by sex if significant differences were found; otherwise, mixed-sex cohort data are shown. Additional sex-separated data are in supplemental figures 2 and 3. For 1-way or Welch ANOVAs, Holm’s-Sidak (HS), Tukey’s, or Dunnett’s T3 (D.T3) post-hoc tests compared all experimental groups. For 2-way ANOVAs, HS tests compared groups within each variable (e.g., genotype or sex). Experimental n’s represent individual mice. Significance was defined as *p <*0.05 with markings * p<0.05, ** p<0.01 *** p<0.001, and **** p<0.0001. Data are expressed as the mean ± SEM.

## DATA AVAILABILITY

The datasets used and/or analyzed during the current study are available from the corresponding author on reasonable request.

## ACKNOWLEDGEMENTS/AUTHOR CONTRIBUTIONS

JH conceptualized and designed the study. AH conducted open field and grip strength assays. NS carried out the marble burying, social experiments, and genotyping. AH and NS completed rotarod and gait studies. Nesting was performed by JH and JR. JR completed the PPI. The fear conditioning experiment was conducted by JH, AH, and JR. JH conducted the ataxia and glucose assays. SM, AH, and JH collected and analyzed H-reflex measurements. Auditory assessments were conducted by NS and AH, with data analysis completed by PW, JH, and AH. JGu provided the mice and contributed to data interpretation. NS, AH, JR, JH, JGr collected end-point tissue. Golgi staining was performed by JR and NS, followed by imaging and analysis by JR. JGr conducted the western blots. JH supervised and assisted in all data analysis. JH and JGr wrote the manuscript, which was reviewed and approved by all authors.

## DECLARATION OF GENERATIVE AI AND AI-ASSISTED TECHNOLOGIES IN THE MANUSCRIPT PREPARATION PROCESS

During the preparation of this work the authors used UR AI and ChatGPT to improve readability and language. After using this service, the authors reviewed and edited the content as needed and take full responsibility for the content of the published article.

