## Supplemental for "Loss of SEZ6L2 disrupts synaptic architecture and drives progressive behavioral deficits"

**Supplemental Figures**

Sup_Fig 1

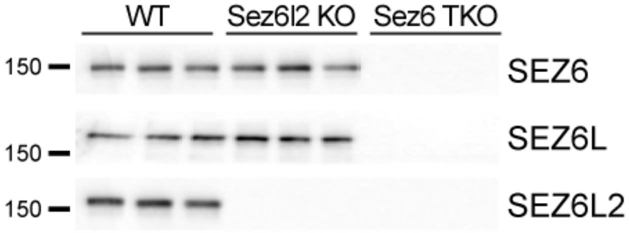


**Supplemental Figure 1: Validation of mouse strains.** Western blot of cortical lysates from WT, *Sez6l2* KO, and *Sez6* TKO mice blotted for SEZ6, SEZ6L, and SEZ6L2.

Sup_Fig 2

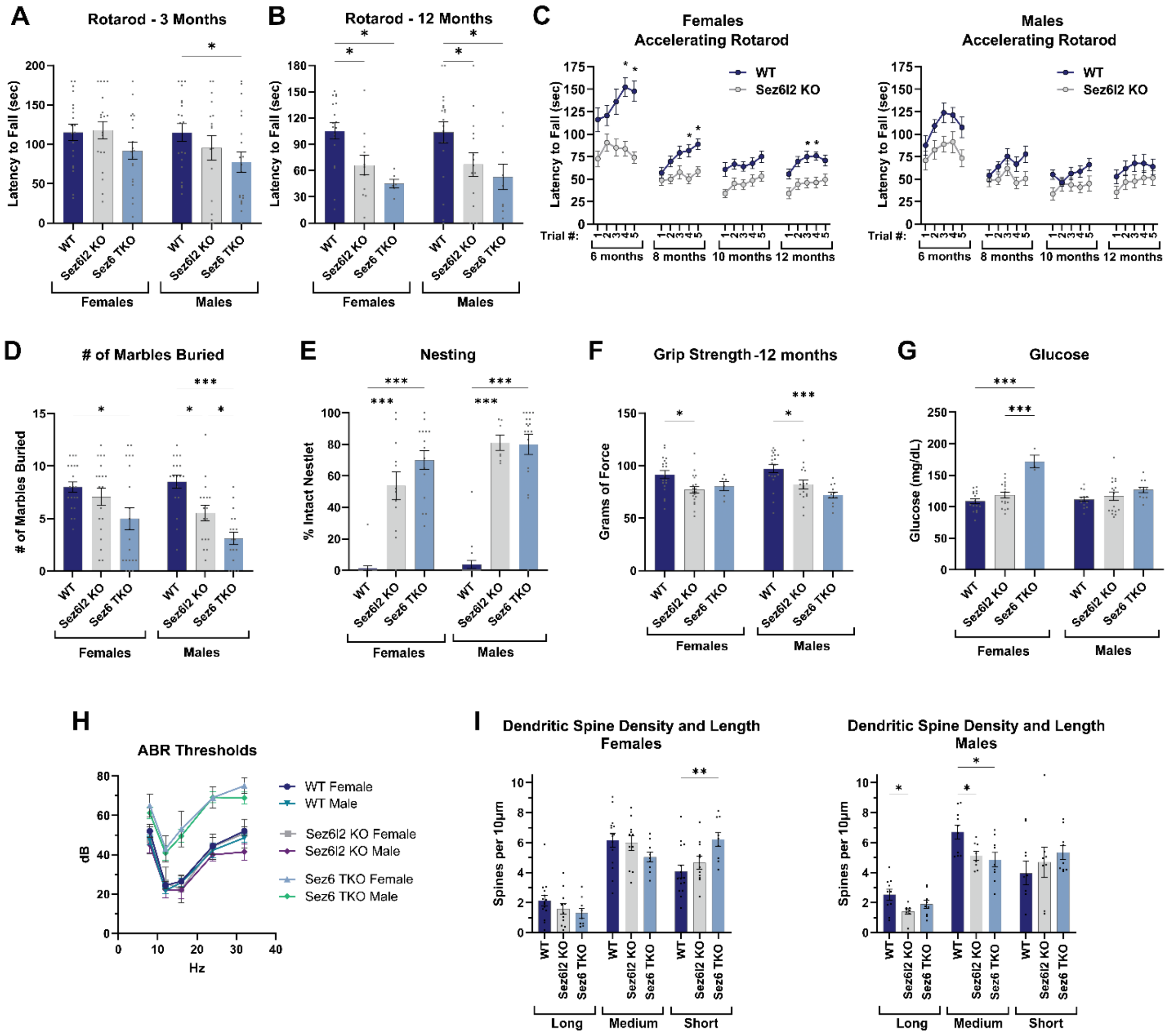


**Supplemental Figure 2: Select Male and Female Data. A,B)** Latency to fall on fixed speed rotarod (12 RPM) at 3 and 12 months3 months: WT N=40 (20F, 20M); *Sez6l2* KO N=36 (19F, 17M); *Sez6* TKO N=39 (20F, 19M); 12 months: WT n=37 (18F, 19M); *Sez6*L2 KO n=27 (12F, 15M); *Sez6* TKO n=15 (6F, 9M). **C)** Female and male WT and *Sez6l2* KO mice were evaluated for latency to fall on an accelerating rotarod at 6, 8, 10, and 12 months with 5 trials per timepoint. For all time points: WT n=38 (19F, 19M); *Sez6l2* KO n=39 (22F, 17M). **D)** Number of marbles buried in

30 minutes due to repetitive digging. WT n=40 (20F, 20M); *Sez6l2* KO n=36 (19F, 17M); *Sez6* TKO n=39 (20F, 19M). **E)** Nesting assay; percent of nestlet that remained intact after 24 hours. WT n=41 (21F, 20M); *Sez6l2* KO n=19 (12F, 7M); *Sez6* TKO n=33 (15F, 18M). **F)** Quantification of forelimb grip strength at 12 months. WT n=38 (19F, 19M); *Sez6l2* KO n=36 (19F, 17M); *Sez6* TKO n=18 (6F, 12M). **G)** Fasting Glucose measured at 12 months. WT n=30 (16F, 14M), *Sez6l2* KO n=34 (17F, 17M), *Sez6* TKO n=14 (3F, 11M). **H)** ABR thresholds; WT n=21 (10F, 11M); *Sez6l2* KO n=13 (6F, 7M); *Sez6* TKO n=17 (5F, 12M). **I)** Quantification of spine density binned by length. WT n=25 dendrites (15F, 10M); *Sez6l2* KO n=19 (11F, 8M); *Sez6* TKO n=18 (9F, 9M). Statistics for all graphs: 2-Way ANOVAs with HS MCTs.

Sup_Fig 3

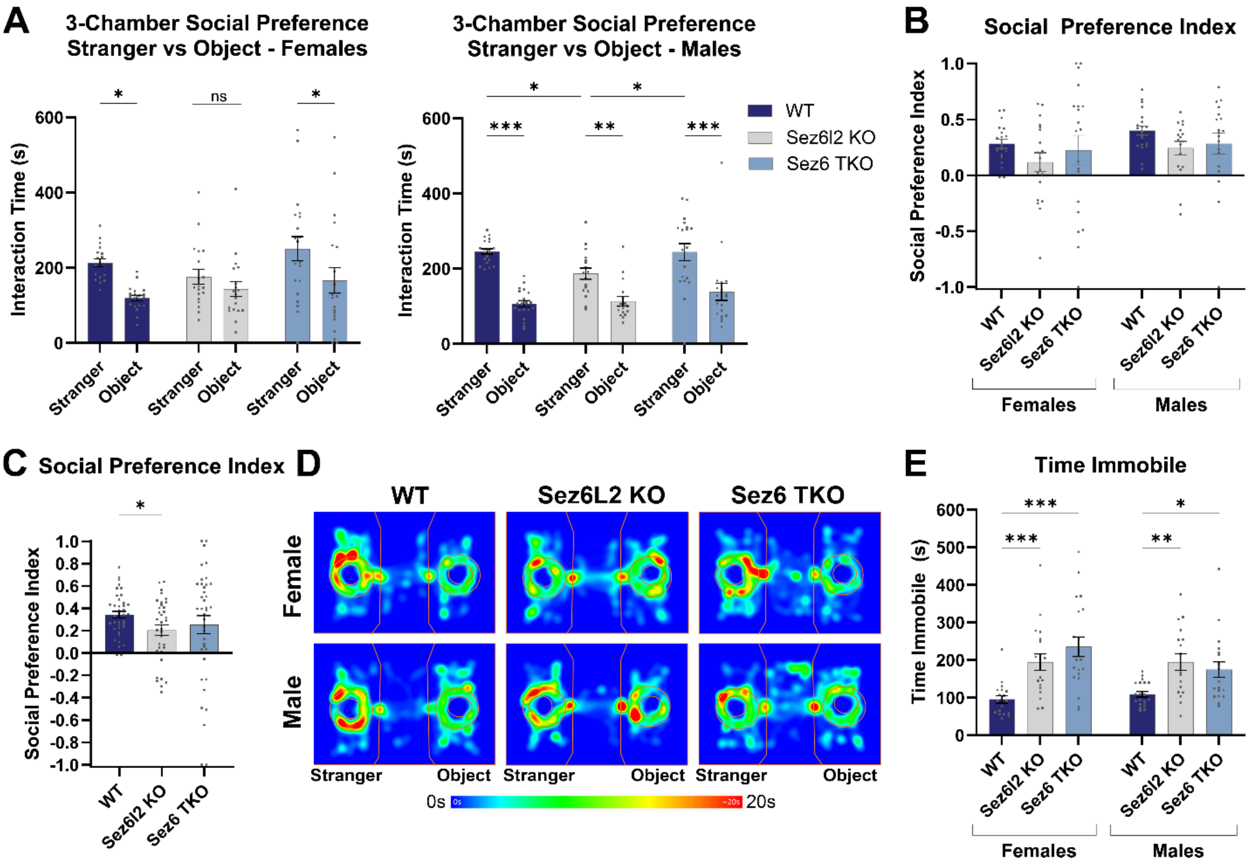


**Supplemental Figure 3: Male and Female Data in 3-chamber stranger versus object test. A)** Quantification of the time female and male mice spent interacting with the stranger mouse versus the object. **B, C)** Bar graphs comparing the social preference indexes of each genotype separated by sex (B) or as a mixed-sex cohort (C). D**)** Representative heatmaps displaying the spatial and time patterns of each genotype in the social preference test. **E)** Quantification of the time mice of each genotype were immobile during the social preference test. Statistics for graphs in A: 2-way RM ANOVA with HS MCTs. B and E: 2-way ANOVA with HS MCTs; and C: Welch ANOVAs with D.T3 MCT. For all graphs: WT n=40 (20F, 20M); *Sez6l2* KO n=36 (19F, 17M); *Sez6* TKO n=40 (20F, 19M).
